# Evolvoid: A genetic algorithm for shaping optimal cellular constructs

**DOI:** 10.1101/2024.09.24.614676

**Authors:** P. Mancini, F. Fontana, E. Botte, C. Magliaro, A. Ahluwalia

## Abstract

We describe an *in-silico* pipeline, Evolvoid, based on Genetic Algorithms (GAs) for identifying the optimal morphologies of cell-laden constructs. Driven by an ad hoc selection rule (i.e., the so-called fitness function (FF)), Evolvoid iteratively identifies the characteristics (i.e., the genome) of the ‘survival of the fittest’ individual of a given population throughout generations. The FF is based on universally observed biophysical laws, representing the optimal trade-off between *i)* high cell viability and robustness to changes in environmental oxygen and *ii)* a low surface energy. The Shannon entropy is used to evaluate genome complexity, with the most complex fittest individuals showing quantitative and qualitative biological resemblance to *in vitro* constructs. Evolvoid paves the way for the development of “*lab on a laptop*”: high-fidelity and cost-effective digital twins of cellular constructs which could augment or even substitute costly *in vitro* models.

## Introduction

Morphogenesis (*i*.*e*., the emergence of shape in living organisms) is among the most complex, multiscale processes observable in nature. The investigation and modelling of morphogenesis has gathered increasing interest as an aid to interpreting the structure-function relationship of biological systems and understanding how cells might cooperate to deal with perturbations to generate reproducible morphologies [1], [2], [3]. Several studies have shown that cell differentiation and aggregation during development is driven by several biophysical constraints – such as minimization of surface energy and adequate nutrient supply – which define organismal shape according to universal laws[4].

In principle, the same constraints should also hold for cells which self-aggregate *in vitro* to form three-dimensional (3D) constructs (*e*.*g*., spheroids, organoids). Therefore, the prediction of these phenomena could enhance the fidelity of their digital twins, which represent a valuable tool to optimize experimental practice. However, besides a few examples of agent-based models[5], digital models of 3D cell constructs currently rely on deterministic and simplistic shapes-such as perfect spheres or ellipses [6] – which are far from the irregular morphologies observed experimentally [7]. This assumption could impair the reliability of computational simulations.

Morphogenesis is driven by genetic programs, intercellular interactions and microenvironmental cues. Indeed, several reports deal with morphogenesis from an evolutionary perspective, studying form as a consequence of life-history optimization [8]. Biophysical constraints are known to impact cell behaviour by modulating cell shape, which can also directly drive the morphogenesis of tissues. From a mathematical point of view, evolution can be considered as a sophisticated optimization process, enabling living systems to achieve highly functional solutions to address specific environmental needs. This insight has inspired novel approaches for numerical optimization through what is referred to as evolutionary computation (EC) [9], which implements the concepts of Darwinian evolution to solve metaheuristic problems [10]. *Genetic algorithms* (GAs) represent the best-known EC method and are governed by the “survival of the fittest” law [11]. Specifically, they identify the characteristics (*i*.*e*., the genome) of the optimal solution (*i*.*e*., the fittest individual of the population) iteratively throughout generations. The optimal solution minimizes/maximizes a predefined objective function (*i*.*e*., the so-called *fitness function*), which sets the rules of the selection process. In this regard, Kriegman and coworkers[12] have coupled GAs with artificial intelligence to develop a pipeline for designing reconfigurable organisms able to complete specific locomotion tasks. The authors artificially crafted task-oriented biobots (*i*.*e*., robots made up of biological material) through surgical manipulation of living tissues, offering a powerful toolkit to fulfil complex functions in hardly accessible contexts.

Here, we use a complementary approach; rather than optimization of locomotory performance for generating new forms, the selection rules are based on biophysical laws which drive the evolution of living systems. Based on studies of spontaneous cell aggregation *in vitro*, we use GAs to identify the optimal shape of randomly generated aggregates resembling salient structural and functional features of *in vivo* tissue: i) three-dimensionality, ii) the minimization of surface energy, iii) maintenance of viability and robustness to perturbations in environmental O_2_ [13]. The GAs are embedded in a computational pipeline named Evolvoid (Fig. 1), which relies on the integration of evolutionary and finite element (FE) computation with the functional information theory of evolution[14]. Specifically, the intrinsic properties of evolutive processes in terms of their complexity and function are leveraged as quantitative benchmarks for assessing the biological relevance of outcoming cell aggregates, through an ad-hoc fitness function (FF). Starting from populations of fifty individuals (see SI 1) with the same average core radius (R) and random geometrical traits (*i*.*e*., morphological genes, Fig. 1, step 0), the pipeline determines optimal shapes complying with biophysical constraints. The identification of the biophysical genes (Fig. 1, Step 2)-which carry information about cell viability, surface energy and three-dimensionality-is accomplished by simulating individual dynamics through FE models. Subsequently, individuals are scored through the FF (Fig. 1, Step 3) determining the next generations of individuals (i.e., the offspring), composed of the elite (i.e., replicates of parents with the highest FF values) and recombinant individuals (Fig. 1, Step 4). The best-scored and hence optimal construct according to the FF-is identified. The operations are run iteratively within the GA cycle until predefined stop conditions are met.

**Fig 1.**
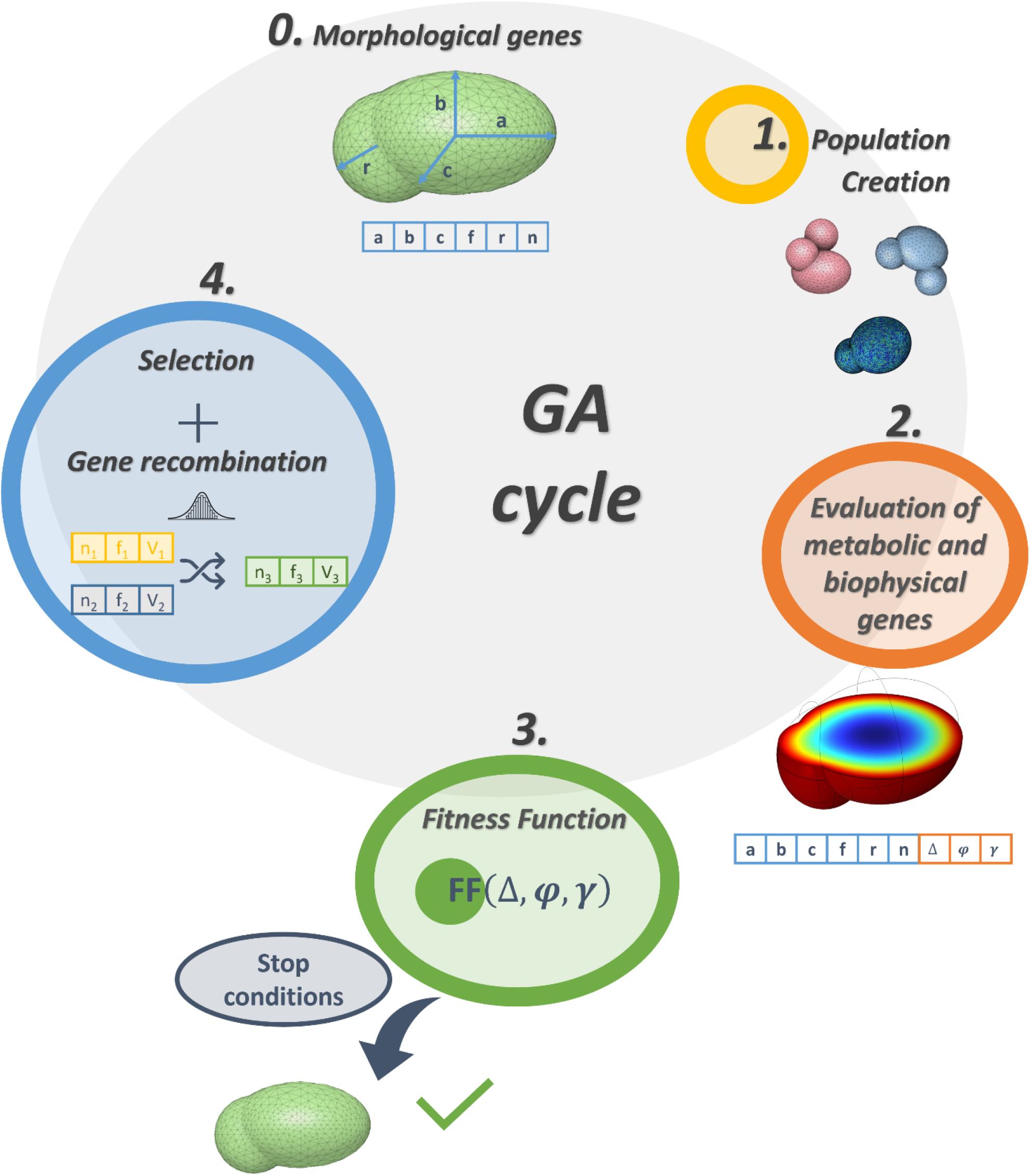
Evolvoid’s pipeline: Morphological genes (0) are selected to generate a population of constructs (1). The optimal macroscale morphology of the constructs is determined iteratively by evaluating biophysical genes (2). Each individual is thus associated with its genome, which contains a set of morphological and biophysical genes. The fitness function, FF is evaluated (3) and drives the generation of successive populations through selection and gene recombination (4). The GA cycle iteratively computes until predefined stop conditions are met. In this way, the best scored and hence optimal construct according to the FF is identified.

As a first assessment of Evolvoid’s performance, focusing on hepatic constructs, we ran the GA cycle for different starting population average radii (R), comparing the optimal shapes from the pipeline with their *in vitro* counterparts. Albeit based on a limited number of fundamental selection rules, the fittest morphologies are remarkably similar in both shape and size to experimental constructs reported in the literature. Evolvoid thus provides the basis for a new lap on a laptop-based methodology for refining, reducing and replacing both *in vitro* and animal experiments in biomedical research. It paves the way for the generation of high-fidelity digital twins of cell-based models, enabling the optimization of their design for applications such as tissue engineering, drug and biomaterial testing or the investigation of specific dynamics that underpin emergent biological behaviours.

## Results

Figure 2 schematises the results of the pipeline. In particular, starting from a random population and based on a FF which quantifies morphological and biophysical descriptors, the optimal shape is identified across generations and compared with its *in vitro* counterpart (Fig. 2A). The optimal individual for a core radius *R* = 521μm along with its genome, is shown in Figure 2B. As illustrated in the figure, the genome is composed of the morphological (*a, b, c, r, n, f*) and biophysical genes (*Δ, φ, γ*). This genome has a high FF value across generations, indicative of an optimal trade-off between *i)* high cell viability and robustness to changes in environmental oxygen (*i*.*e*., the change in viable volume fraction *Δ*) and *ii)* a low surface energy by maximizing both sphericity (*φ*) [15] and three dimensionality (*i*.*e*., surface area to volume ratio of the construct with respect to that of a sphere of radius *R, γ*. See Section A2, Table 2).

**Fig 2.**
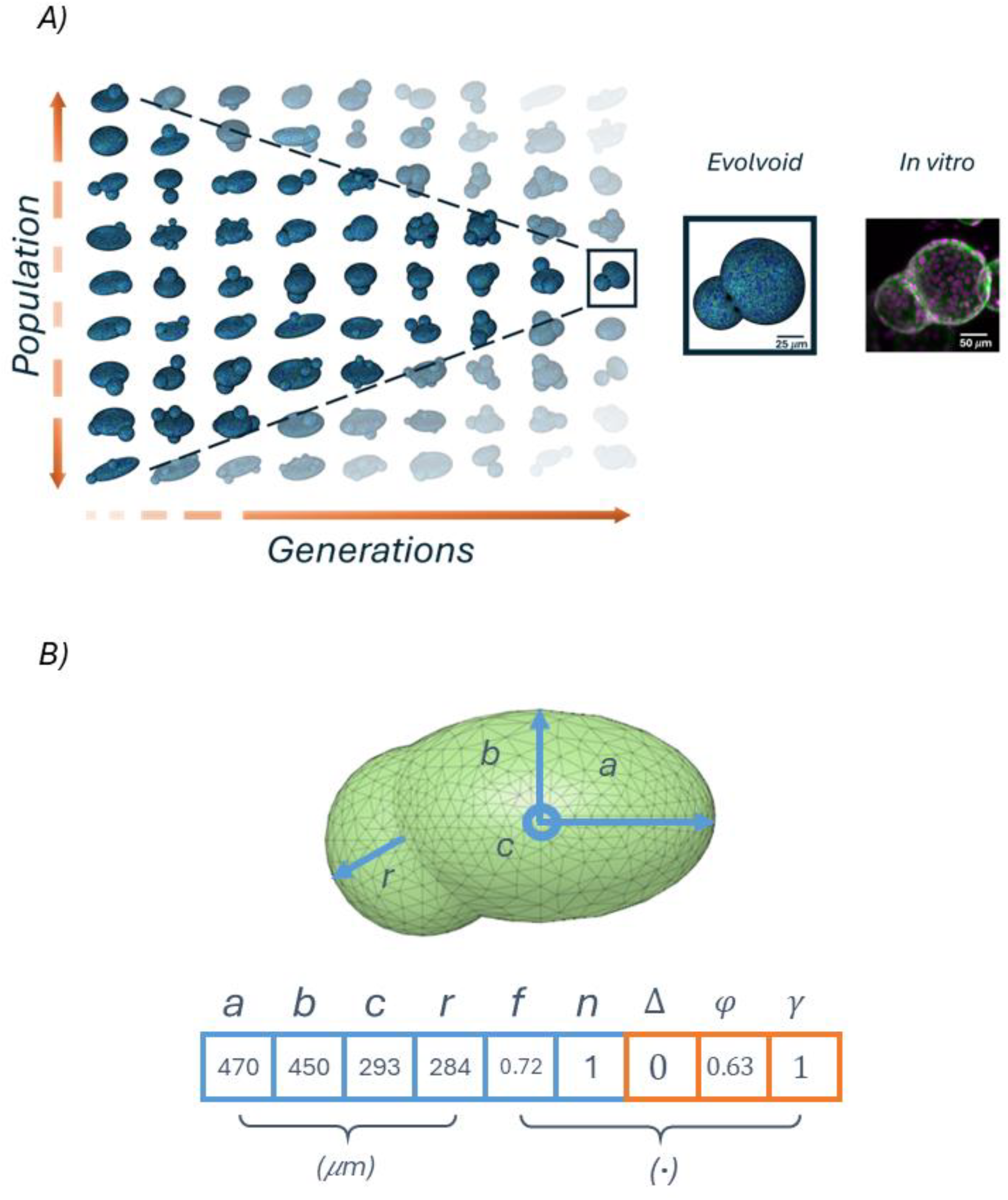
A) Outcome of the pipeline showing a subset of construct populations and how the process of selection and recombination leads to the emergence of optimal morphologies through generations. B)Example of an optimal individual and its genome, comprising both morphological and biophysical genes. The morphological genes define the semiaxes of the construct core (a, b, c), the radius of protrusions 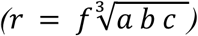, the number of protrusions (n), and the number of protrusions-to-central ellipsoid ratio 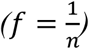 (See Section A1). The biophysical genes (Δ, φ, γ) express the change in viable volume fraction in the presence of two different oxygen boundary levels, the sphericity [15] and the three dimensionality, respectively (Section A2).

A typical trend of the FF evaluated across generations expressed in terms of mean ± standard deviation per generation is plotted in Figure 3A: the bounded region is magnified pointing out three individuals (Fig. 2B), with the highest (individual n°1), the mean and the lowest (individuals n° 2 and n° 3, respectively) FF values for their respective generation. In fact, the constructs clearly exhibit different morphologies (Fig 2C). More in detail, the optimal individual (n°1) displays the best combination of the number of protrusions, their size (*f* = 0.72), and their interpenetration with the core (see Section A and SI 2B), ensuring the minimization of the surface energy while maintaining overall cell viability robust to environmental oxygen concentration. On the other hand, suboptimal individuals show a flatter core (Fig. 3C, individual n° 3) or a higher surface area with a high number of protrusions (Fig. 3C, individual n°2). Although both these morphologies promote nutrient supply and thus have a high Δ, they have a lower overall FF than the one calculated for the optimum. The flat individual has a low three dimensionality, *γ*, while the construct with a high number of protrusions has low sphericity *φ*. Figure 3D shows how the maximum value of FF increases as a function of generation leading to the identification of the optimal individual due to stop conditions (see SI 3 for further details).

**Fig 3.**
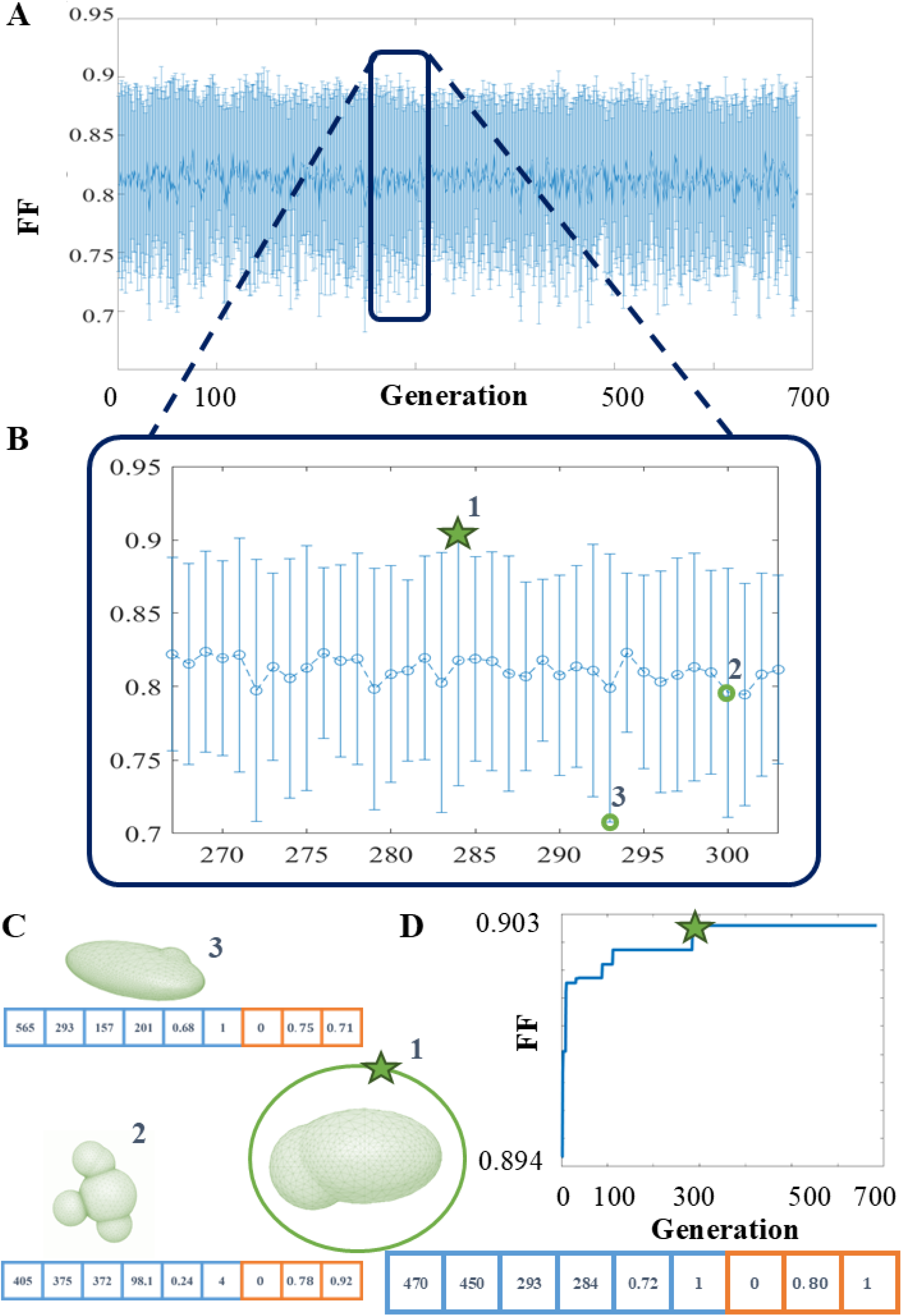
Evolvoid, a typical outcome. **A)** Mean ± standard deviation of the FF for the 50 individuals of each generation as they evolve over 400 GA cycles. Upper and lower limits correspond to the best- and worst-fitting individuals of each generation, respectively. B) The trend in more detail for about 30 cycles (generations 270 to 305). **C)** The individual n° 1 is the optimum with the highest FF, followed by the n°2 and then n°3. **D)** Trend of the maximum value of the FF across generations. The global maximum of the FF is reached when the optimal individual first appears, i.e., at the 227^th^ generation. The search process stops after the stop conditions are met (SI 3).

**Fig 4.**
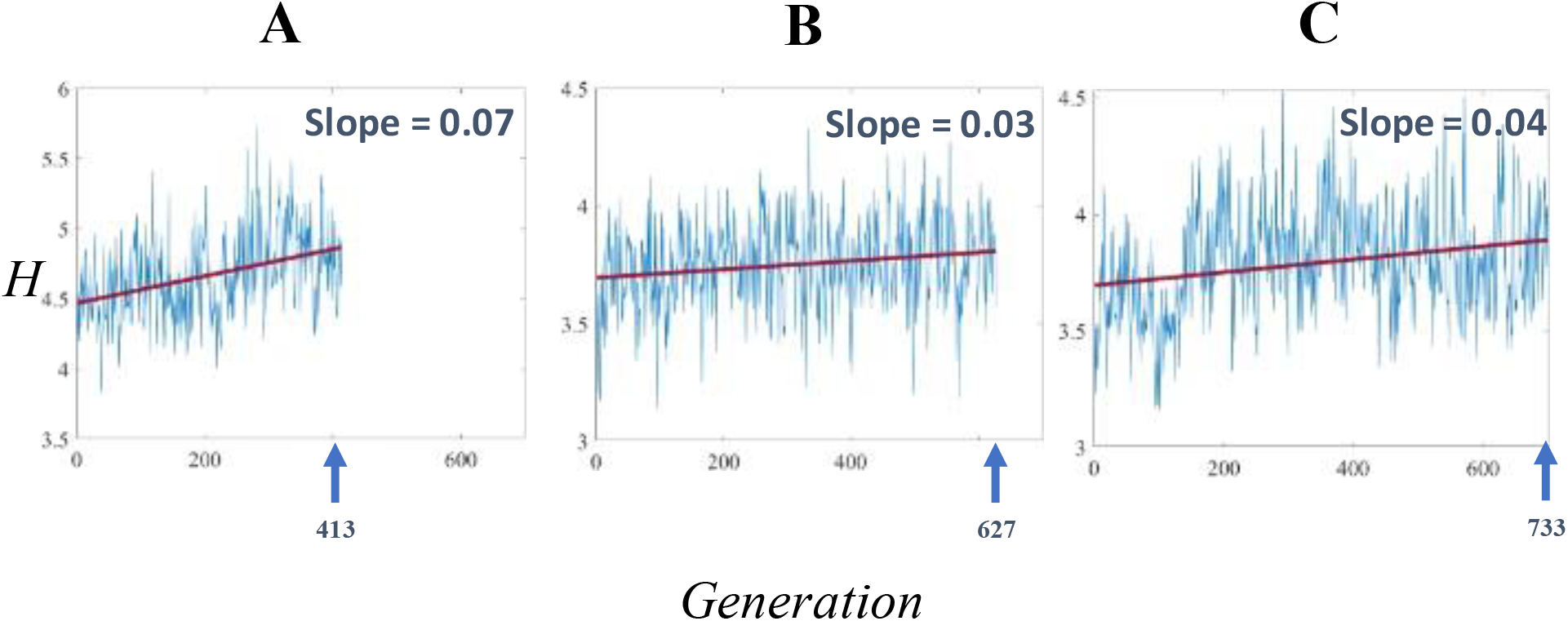
Construct complexity H across generations with the associated linear fitting (in red) and the corresponding slope value, of three runs of the algorithm with different starting average core radii R. Non-parametric Spearman correlation coefficient (Sp) and the associated p-value referring to the null hypothesis that the two variables are uncorrelated (p_r_) are evaluated. A) R=654 μm, p-value of the slope significantly non-zero (p_s_)< 0.0001, Sp=0.16, p_c_ = 0.0099. B) R= 521 μm, p_s_<0.0001, Sp=0.39, p_c_ <0.0001. C) R= 918 μm, p_s_<0.0001, Sp=0.25, p_c_ < 0.0001.

**Fig 5.**
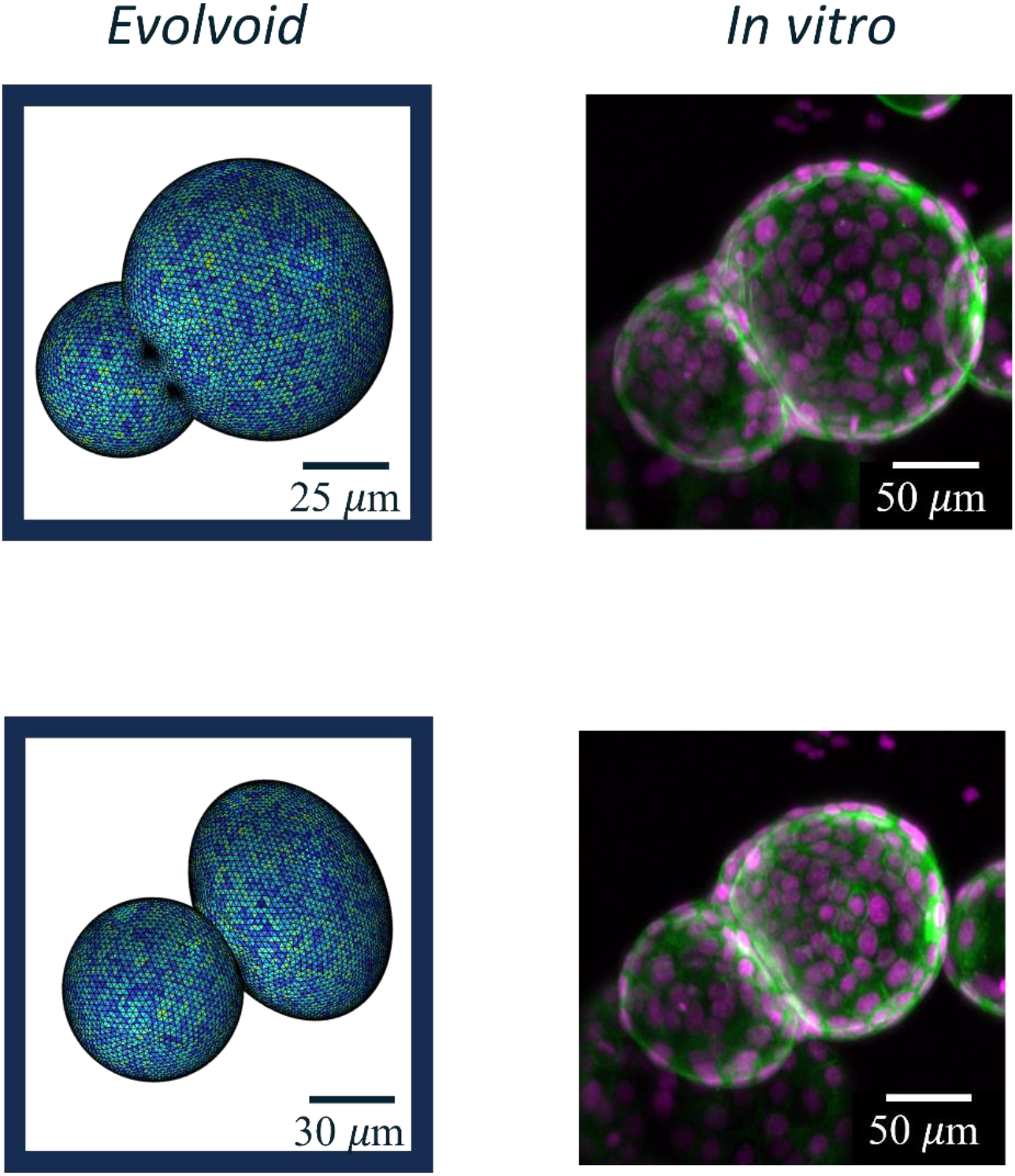
Comparison of optimal constructs obtained with Evolvoid (left) and images of human cholangiocarcinoma-derived organoids (hCCAOs, right)[7].

### Genomic complexity

In Evolvoid, the construct complexity-computed as the Shannon entropy, H-quantifies the intra-generation variability of individuals (See Section D and SI 4). Due to the stochasticity of the search process, H, across generations, is a fluctuating signal. For this reason, a linear fitting is used for visualizing the trend of the average complexity of the process. In coherence with all evolutive processes in nature[16], a gradual increase in construct complexity across generations is observed. The slope of the linear fitting positively correlates with the number of generations (Spearman correlation coefficient, Sp > 0 with *p*_*c*_ < 0.01).

### In vitro comparison

The optimal constructs returned by Evolvoid have sizes comparable to those experimentally observed *in vitro*. We compared images of human cholangiocarcinoma-derived organoids (hCCAOs) in suspension[7] with the optimal morphologies predicted by Evolvoid. The organoids are qualitatively similar in shape and size as shown in Figure 5. Furthermore, the optimal constructs generated by Evolvoid with core radius *R*_1_ = 31 *um* and *R*_*2*_ = 54 *um*, display a size range comparable to that of HepG2 spheroids obtained with hanging-drop method[17].

## Discussion

Evolvoid is a computational platform based on GAs for reliably predicting the shape and size of cellular constructs. It relies on the maximization of a FF which balances the morphological features of each construct with relevant biophysical criteria [13], [18], [19], [20], so as to identify the best-fitting geometrical and viability traits [15], [21], [22]. Although the FF needs further enhancement by means of incorporating additional biophysical constraints (*e*.*g*., the minimization of elastic energy within the construct [23] and compliance with scaling laws [24], [25]) for widening the space of possible solutions [10], [11], [12] and therefore, improving the model predictivity, the results of this study highlight that Evolvoid generates morphologies with resemblance to 3D cell constructs generated in *in vitro* experiments. Leveraging the analogy between Shannon entropy and genomic complexity for the intra-generation variability of individuals, we demonstrate that morphologies generated through Evolvoid’s GA cycles undergo a generic increase in complexity with a statistically significant non-zero slope, aligning with evolutionary processes. Given its computational architecture, Evolvoid can span a wide spectrum of optimal constructs in terms of both sizes and cell types involved, by tuning the equivalent core radius (R, which *in vitro* is essentially constrained by the fabrication technique, including cell number and well-size) and the values of the kinetic parameters for O_2_ consumption, respectively. More generally, the modularity of Evolvoid opens the possibility to model various *in vitro* constructs and develop ad hoc *in silico* models. In fact, this can be achieved by adjusting the FF and thus, driving evolution in specific ways, by modifying the number of genes to increase the individual’s complexity and search space, and by integrating the FE model (e.g., introducing additional consumed species or considering a time-dependent model). Evolvoid paves the way for improving digital twins of cell-based *in vitro* models and for developing the lab-on-a-laptop concept with an even more high throughput technology and represents a groundbreaking advancement in the development of new approach methodologies (NAMs) for refining, reducing and replacing both *in vitro* and animal models in biomedical research.

## Methods

### A. Generation of populations

Using the pipeline shown in Figure 1, fifty individuals sharing the same average core radius R, were generated. For each run, R was randomly picked from one of fourteen lognormal distributions with means ranging from [31 um, 2142 um], as in [24], [25]. The geometrical features of the individuals constitute their morphological genes, which are then used to compute their biophysical genes through FEM analysis.

#### A.1 Morphological genes

In the most general description of construct morphology, individuals have an ellipsoidal core (i.e., they are slightly flattened since gravitational forces predominate over surface tension [26]), with smaller spherical protrusions (with lower Bond number) randomly positioned on the surface (see SI 2B). The three-semiaxes of the ellipsoid are calculated as a fraction of *R* (see SI 2A), picked from a uniform distribution (*U*_*R*_) ranging from [0, 1]. On the other hand, the size of protrusions (*r*), and their number (*n*) are related to a size fraction (*f*). The size fraction is inversely proportional to *n*, because of steric hindrance and scaling and resource allocation requirements [27], [28]. Table 1 and Figure 3A summarize all the morphological genes of an individual. Further details are given in SI 2.

**Table 1.**
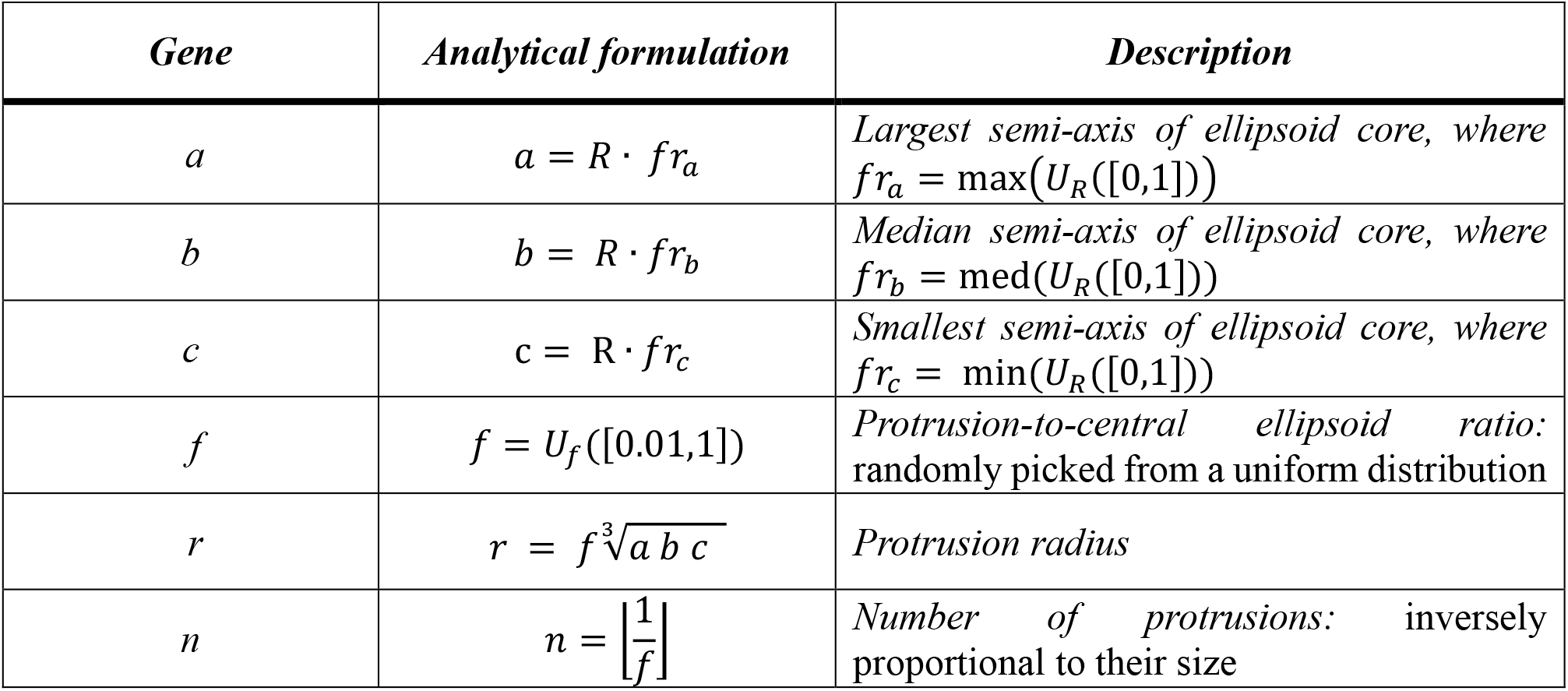
Morphological genes considered for determining shape and size of individuals in Evolvoid. U([x, y]) signifies a uniform distribution within the range x:y and R is the core radius.

**Table 2.**
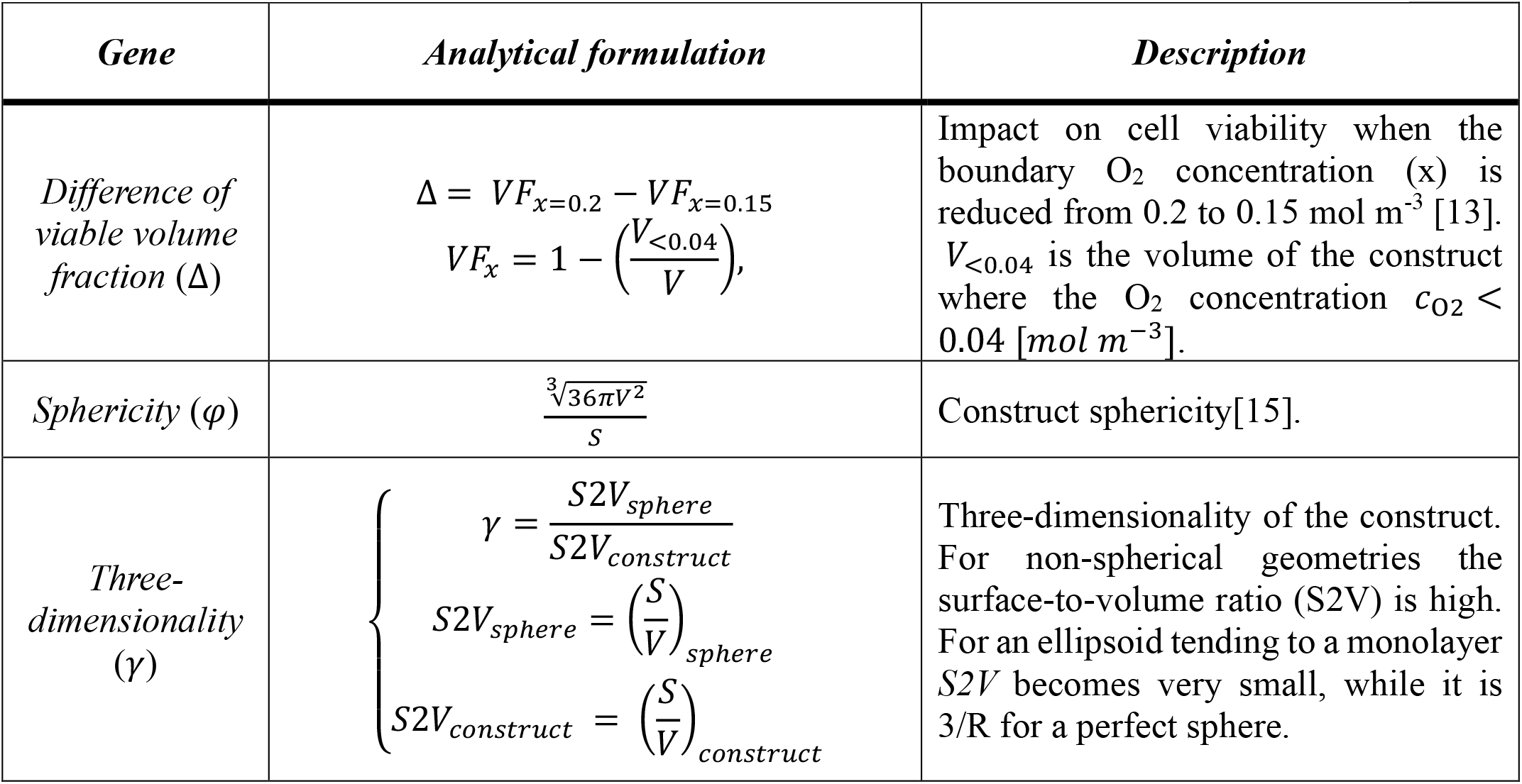
Biophysical genes considered as criteria for the selection process in Evolvoid. V [m^3^] is the total volume of the construct and S [m^2^] is its total surface area.

#### A.2 Biophysical genes

Once all morphological genes describing the geometry of each construct have been assigned, O_2_ transport and consumption are modelled through FE analysis using the reaction-diffusion equation:

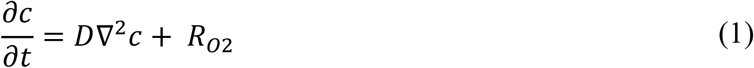

In Eq. (1), *c* (mol m^-3^) is the O_2_ concentration, *D* (m^2^ s^-1^) the diffusion coefficient of O_2_ in water at 37°C and *R*_*O2*_ (mol m^-3^ s^-1^) the O_2_ consumption rate, which typically follows Michaelis-Menten (MM) kinetics[29].

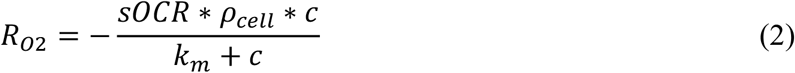

Eq. (2) assumes a construct with a uniform cell density, *ρ*_*cell*_ (cell m^-3^), while *sOCR* (mol s^-1^) and *km* (mol m^-3^) denote the single cell O_2_ consumption rate and the MM constant, respectively. Eq (1) was solved using FE analysis to compute the oxygen concentration field at steady state and with different boundary conditions, thus defining the biophysical genes (Table 2). The models were implemented in COMSOL Multiphysics (version 6.0, Stockholm, Sweden) combined with Matlab (R2022a, MathWorks, Massachusetts, USA) through COMSOL LiveLink™ for Matlab (SI 5).

All parameter values and boundary conditions considered are listed in Table SI 1.

### B. Fitness function

GAs require an optimization function, the so called fitness function (FF), which describes a quantitative benchmark for finding the best solution to a given problem. The choice of the FF is essential for implementing a well-posed and effectively converging search process, since it defines the space of possible solutions explored by the GA. In our case, the aim is to identify the best shaped individuals, thus the FF should tend to a single, global optimum solution [19], [20].

The FF was implemented as a damped harmonic oscillator (Eq. (3)); this dynamic behaviour is observed in several biological processes (*e*.*g*., circadian rhythms)[30], [31], [32]. The damping factor (*wΔ*) guarantees that, on average, cell constructs with a higher difference of viable volume fraction, Δ, are associated with a lower score, due to their limited resistance to environmental perturbations. On the other hand, the genes associated with sphericity, *φ*, and three-dimensionality, *γ*, are introduced in the score by means of a Lagrange multiplier, *w*_<*φ,γ*>_ (typical of constrained optimization problems) [33]. See the SI6 for further details.

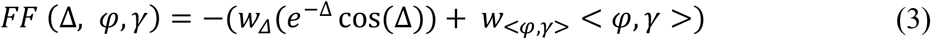

### C. Successive generations

The FF is used to score all the individuals of a given population, to generate offspring. The best-scored (5%) of them became the so-called *elite* (*i*.*e*., closest to the optimum). Non-elite individuals (NEIs) generate offspring through gene recombination: *crossover* and *mutation*. Crossovered genomes are obtained by combining the morphological genes of two different NEIs, while the mutated ones are randomly generated modifying one (or more) morphological gene(s) from a single NEI. Crossover allows densifying the search space, since it generates individuals owning an already existing morphological features which are combined differently. On the other hand, random mutation increases the diversity of the population, enabling the exploration of a wider spectrum of possible solutions. The fraction of NEIs undergoing crossover is randomly set according to a gamma probability density function (as specified in SI7-Table SI2)[34], and the fraction of mutated progeny is derived consequently. Once the population is completed, the GA cycle is run iteratively until a stop condition (SI 3) is verified.

### D. Analysis of genomic complexity

The complexity of a biological system increases as it evolves [16]. If considered as a string of characters which are randomly combined and compared with a target string, the process of evolution can typically be evaluated in terms of information theory entropy [21], [22]. In fact, given a formal analogy with information theory[14], the complexity of an individual generated with Evolvoid and thus, the total complexity of the generation, can be quantified as the Shannon entropy of the morphological genes, which sums all the possible values the genes could have (See Eq. (S15) in SI 4). The biological relevance of Evolvoid is thus assessed by determining the average morphological complexity of individuals across generations through Eq. (S15) in SI 4. The formula and the definitions used are detailed in SI 4. GraphPad Prism (version 7, GraphPad Software, California, USA) was used for statistical analysis.

Further details on the finite element computation and data analysis are provided in SI 5.

## Supporting information

Further detailes on the computational implementation of Evolvoid

## Acknowledgments

This work received funding from the Swiss National Science Foundation, through Project SINERGIA 2019, under Grant Agreement No. 572 CRSII5_186422/1. EB’s post-doctoral fellowship was funded by the Next Generation EU Project CN00000013—”Centro Nazionale 1:” HPC, Big Data and Quantum Computing (CN1, PNRR, Spoke 6: Multiscale modelling and engineering applications). This research has received funding from the Italian Ministry of Health for the development of alternatives to animal models.

## Author declarations

### Ethics Approval

Ethics approval is not required.

## Author Contributions

Piera Mancini and Flavio Fontana contributed equally to this work.

*Piera Mancini*: Conceptualization (equal); Data curation (equal); Formal analysis (equal); Investigation (equal); Methodology (equal); Software (equal); Writing – original draft (equal); Writing – review & editing (equal).

*Flavio Fontana*: Conceptualization (equal); Data curation (equal); Formal analysis (equal); Investigation (lead); Methodology (equal); Software (equal); Writing – original draft (supporting); Writing – review & editing (equal).

*Ermes Botte:* Conceptualization (supporting); Data curation (equal); Formal analysis (equal); Investigation (equal); Methodology (supporting); Software (supporting); Supervision (supporting); Writing – original draft (supporting); Writing – review & editing (equal).

*Chiara Magliaro*: Conceptualization (lead); Data curation (equal); Formal analysis (equal); Investigation (equal); Methodology (equal); Software (equal); Supervision (lead); Writing – original draft (supporting); Writing – review & editing (equal).

*Arti Devi Ahluwalia*: Conceptualization (lead); Data curation (equal); Formal analysis (lead); Funding acquisition (lead); Investigation (lead); Methodology (equal); Supervision (lead);Writing – review & editing (lead).

## References

[1] A. M. Turing, ‘The Chemical Basis of Morphogenesis’, Philos Trans R Soc Lond B Biol Sci, vol. 237, no. 641, pp. 37–72, 1952, [Online]. Available: http://www.jstor.org/about/terms.html.

[2] M. Epstein and G. A. Maugin, ‘Thermomechanics of volumetric growth in uniform bodies’, Int J Plast, vol. 16, no. 7, pp. 951–978, Jun. 2000, doi: 10.1016/S0749-6419(99)00081-9.

[3] L. Wolpert, C. Tickle, and A. M. Arias, ‘Principles of Development’, Oxford University Press, pp. 1–695, 2015, [Online]. Available: https://books.google.co.in/books?id=dfvVoQEACAAJ

[4] L. Wolpert, ‘Positional Information and the Spatial Pattern of Cellular Differentiationt’, Theoret. Biol, vol. 25, pp. 1–47, 1969.

[5] D. Oriola et al., ‘Arrested coalescence of multicellular aggregates’, Soft Matter, vol. 18, p. 3771, 2022, doi: 10.1039/d2sm00063f.

[6] S. Montes-Olivas, L. Marucci, and M. Homer, ‘Mathematical Models of Organoid Cultures’, Front Genet, vol. 10, p. 465709, Sep. 2019, doi: 10.3389/FGENE.2019.00873/BIBTEX.

[7] L. Hof et al., ‘Long-term live imaging and multiscale analysis identify heterogeneity and core principles of epithelial organoid morphogenesis’, BMC Biol, vol. 19, no. 1, pp. 1–22, Dec. 2021, doi: 10.1186/S12915-021-00958-W/FIGURES/7.

[8] V. Balbi and P. Ciarletta, ‘Mathematical Modeling of Morphogenesis in Living Materials’, in Mathematical Models and Methods for Living Systems: Levico Terme, Italy 2014, P. Ciarletta, T. Hillen, H. Othmer, L. Preziosi, D. Trucu, L. Preziosi, M. Chaplain, and A. Pugliese, Eds., Cham: Springer International Publishing, 2016, pp. 211–274. doi: 10.1007/978-3-319-42679-2_4.

[9] K. De Jong, L. Fogel, and H.-P. Schwefel, ‘The Handbook of Evolutionary Computation’, Handbook of Evolutionary Computation, 1997.

[10] C. R. Reeves, J. E. Rowe, and W. B. Langdon, ‘Genetic Algorithms-Principles and Perspectives A Guide to’, Knowledge Engineering Review, vol. 19, no. 2, pp. 185–186, 2004, doi: 10.1017/S026988890423020X.

[11] D. E. Goldberg and J. H. Holland, ‘Genetic Algorithms and Machine Learning’, Mach Learn, vol. 3, no. 2, pp. 95–99, 1988, doi: 10.1023/A:1022602019183.

[12] S. Kriegman, D. Blackiston, M. Levin, and J. Bongard, ‘A scalable pipeline for designing reconfigurable organisms’, PNAS, vol. 117, no. 4, 2020, doi: 10.1073/pnas.1910837117/-/DCSupplemental.

[13] E. Berger et al., ‘Millifluidic culture improves human midbrain organoid vitality and differentiation’, Lab Chip, vol. 18, no. 20, pp. 3172–3183, 2018.

[14] R. M. Hazen, P. L. Griffin, J. M. Carothers, and J. W. Szostak, ‘Functional information and the emergence of biocomplexity’, Proceedings of the National Academy of Sciences, vol. 104, no. suppl_1, pp. 8574–8581, May 2007, doi: 10.1073/PNAS.0701744104.

[15] I. Cruz-Matías et al., ‘Sphericity and roundness computation for particles using the extreme vertices model’, J Comput Sci, vol. 30, pp. 28–40, Jan. 2019, doi: 10.1016/J.JOCS.2018.11.005.

[16] C. Adami, C. Ofria, and T. C. Collier, ‘Evolution of biological complexity’, Proc Natl Acad Sci U S A, vol. 97, no. 9, pp. 4463–4468, Apr. 2000, doi: 10.1073/PNAS.97.9.4463/ASSET/8179608B-046F-4D60-AEBE-FFCF77611CC0/ASSETS/GRAPHIC/PQ0805620004.JPEG.

[17] U.-K. Shah, J. De Oliveira Mallia, N. Singh, K. E. Chapman, S. H. Doak, and G. J. S. Jenkins, ‘A three-dimensional in vitro HepG2 cells liver spheroid model for genotoxicity studies’, Mutat Res Genet Toxicol Environ Mutagen, vol. 825, pp. 51–58, 2018, doi: 10.1016/j.mrgentox.2017.12.005.

[18] M. S. Steinberg, ‘Reconstruction of Tissues by Dissociated Cells’, Science (1979), vol. 141, no. 3579, pp. 401–408, Aug. 1963, doi: 10.1126/science.141.3579.401.

[19] R. A. Foty,’, G. Forgacs, C. M. Pfleger, and M. S. Steinberg’, ‘Liquid Properties of Embryonic Tissues: Measurement of Interfacial Tensions’, American Physical Society, vol. 72, 1994.

[20] G. Forgacs, R. A. Foty, Y. Shafrir, and M. S. Steinberg, ‘Viscoelastic Properties of Living Embryonic Tissues: a Quantitative Study’, Biophys J, vol. 74, pp. 2227–2234, 1998, doi: 10.1016/S0006-3495(98)77932-9.

[21] H. Kantz and T. Schreiber, Nonlinear Time Series Analysis, 2nd ed. Cambridge: Cambridge University Press, 2003. doi: DOI: 10.1017/CBO9780511755798.

[22] Brin, Michael Stuck, and Garrett, ‘Introduction to Dynamical Systems’, Cambridge University Press, 2002.

[23] A. Torres-Sánchez, M. K. Winter, and G. Salbreux, ‘Interacting active surfaces: A model for three-dimensional cell aggregates’, PLoS Comput Biol, vol. 18, no. 12, p. e1010762, Dec. 2022, doi: 10.1371/JOURNAL.PCBI.1010762.

[24] A. Ahluwalia, ‘Allometric scaling in-vitro’, Sci Rep, 2017, doi: 10.1038/srep42113.

[25] E. Botte, F. Biagini, C. Magliaro, A. Rinaldo, A. Maritan, and A. Ahluwalia, ‘Scaling of joint mass and metabolism fluctuations in in silico cell-laden spheroids’, PNAS, 2021, doi: 10.1073/pnas.2025211118/-/DCSupplemental.

[26] E. Sher, T. Bar-Kohany, and A. Rashkovan, ‘Flash-boiling atomization’, Prog Energy Combust Sci, vol. 34, no. 4, pp. 417–439, Aug. 2008, doi: 10.1016/J.PECS.2007.05.001.

[27] Stearns and C. Stephen, ‘The Evolution of Life Histories’, Oxford University Press, pp. 1–249, 1992, doi: 10.1046/J.1420-9101.1993.6020304.X.

[28] A. F. Bennett, ‘Allometry: Scaling. Why Is Animal Size So Important?’, Science (1979), vol. 226, no. 4681, pp. 1412–1413, Dec. 1984, doi: 10.1126/SCIENCE.226.4681.1412.B.

[29] L. Michaelis and M. L. Menten, ‘The kinetics of the inversion effect’, Biochem. Z, vol. 49, pp. 333–369, 1913.

[30] H. De los Santos, E. J. Collins, J. M. Hurley, and K. P. Bennee, ‘Circadian Rhythms in Neurospora Exhibit Biologically Relevant Driven and Damped Harmonic Oscillations’, ACM BCB, pp. 455–463, 2017, doi: 10.1145/3107411.3107420.

[31] S. Koshkin and I. Meyers, ‘Harmonic Oscillators of Mathematical Biology: Many Faces of a Predator-Prey Model’, Mathematics Magazine, vol. 95, no. 3, pp. 172–187, 2022, doi: 10.1080/0025570X.2022.2055424.

[32] T. Ton-That, ‘A generalized mathematical model of biological oscillators’, Cell Mol Biol Lett, vol. 7, no. 1, pp. 101–104, 2002.

[33] A. R. Conn, N. I. M. Gould, and P. L. Toint, ‘Globally convergent augmented Lagrangian algorithm for optimization with general constraints and simple bounds’, SIAM J Numer Anal, vol. 28, no. 2, pp. 545–572, 1991, doi: 10.1137/0728030.

[34] M. S. Mcpeek and T. P. Speed, ‘Modeling Interference in Genetic Recombination’, Genetics, vol. 139, pp. 1031–1044, 1995.

