## Supplementary material for "Evolvoid: A genetic algorithm for shaping optimal cellular constructs": Further detailes on the computational implementation of Evolvoid

### *SI 1: Determination of the Population Size*

Given the six morphological genes involved in the creation of population process, Evolvoid is based on a population size (i.e., numbers of individuals per generation) of fifty according to the considerations in [1] on GAs implementing elitist and roulette wheel selection and the MathWorks guidelines [2]

### *SI 2: Morphological genes*

The morphological genes are the semiaxes of the core $(a, b, c)$, the protrusion radius $(r)$, the protrusion to central ellipsoid ratio $(f)$ and the number of protrusions $(n)$.

#### SI 2.A: Semiaxes of the ellipsoidal core

For the initial run of the algorithm, we establish the first generation of 50 constructs considering an average core radius $(R)$ with the corresponding semiaxes $(a, b, c)$ of the elliptic core. $R$ is randomly selected from a lognormal probability distribution as in [3], [4].

To avoid generating an ellipsoid with very high values of a and b compared with c (tending to a cell monolayer rather than a truly three-dimensional construct), an analytical constraint to the semi axes fraction is imposed. Specifically, we assume that only the surface tension can influence construct shape –since the Bond number is small enough to neglect gravitational effects.

Exploiting the Thomsen approximation for evaluating the surface of an ellipse[5]:

$$S_{ell} = 4\pi \left(\frac{(ab)^p + (ac)^p + (bc)^p}{3}\right)^{\frac{1}{p}} \tag{S1}$$

with p = 1.6075, the surface ($S_{ell}$) to volume ($V_{ell}$) ratio $(StV)$ can be defined as Eq. S2:

$$StV = \frac{S_{ell}}{V_{ell}} \tag{S2}$$

Maximizing the three-dimensionality and neglecting the constant terms, we obtain:

$$\left(\frac{1}{a} + \frac{1}{b} + \frac{1}{c}\right) = \frac{1}{R} \tag{S3}$$

Given the information in Table 2, the formula becomes:

$$\left(\frac{1}{fr_a R}+\frac{1}{fr_b R}+\frac{1}{fr_c R}\right)=\frac{1}{R} \tag{S4}$$

And thus, the semi axes dimensions are constrained and related through:

$$\frac{fr_a fr_b + fr_a fr_c + fr_b fr_c}{fr_a fr_b fr_c}=1 \tag{S5}$$

This ensures a coherent combination of sizes, giving construct with a core having a proportioned aspect ratio[6], and is instrumental for maintaining three-dimensionality.

*SI 2.B: Positioning of the protrusions on the surface of the core*

In organoids and spheroids, protrusive structures often emerge as outgrowths that vary in their degree of integration with the core. According to Steinberg's differential adhesion hypothesis, the extent of this integration—referred to here as compenetration—correlates with the work of adhesion between the core and the protrusion[7]. As this relationship has yet to be fully characterized, we modeled the work of adhesion for each protrusion by its height above the surface of the core (*h*), assigning a random compenetration value to *h* ranging from -*r* to *r*, representing, respectively, full integration within the core to complete detachment.

Specifically, for every protrusion, $h$ is randomly picked from a uniform distribution $U_h([-r,r])$, while its azimuth and elevation with respect to the plane *a, b* are randomly assigned respectively as:

$$\theta = U_\theta([-\pi,\pi])$$

$$\varphi = U_\varphi([-\pi,\pi])$$

The total volume of the construct, considering the protrusion(s), can thus be expressed as:

$$V_{tot} = \frac{4}{3}\pi abc + n\frac{4}{3}\pi r^3 - V_{comp} \tag{S6}$$

Where $abc$ are the three semi-axes, $n$ is the number of protrusions with radius equal to $r$ and $V_{comp}$ is the compenetrating volume (Eq. S7).

$$V_{comp} = \sum_{i=1}^{n}\frac{2}{3}\pi h_i\left(3\sqrt{2h_i r - {h_i}^2} - h_i\right) \tag{S7}$$

Transforming to Cartesian coordinates for computational purposes, the relationships become:

$$\begin{cases} x = (a+h)\cos\theta\cos\varphi \\ y = (b+h)\cos\theta\sin\varphi \\ z = (c+h)\sin\varphi \end{cases} \tag{S8}$$

The number of different values of $h, \theta, \varphi$ for each construct are dependent on the gene $n$ *(number of protrusions)*, which is assigned according to the randomly generated gene *f,* which defines the *protrusion-to-central ellipsoid ratio* (see Table 2, main text). Hence, $h, \theta,$ and $\varphi$

were not inserted in the genome but used rather to explore a wider spectrum of morphologies which might impact on the biophysical genes.

#### *SI 3: Algorithm stopping criterion*

One of the fundamental steps to obtain a well-posed stochastic search process is the definition of an adequate number of points in the search space to obey the infinite population approximation. To achieve this goal, one can work on two aspects:

i. The number of iterations before convergence $t_{max}$;
ii. The number of iterations with the same maximum Fitness Function (FF) value before convergence $t_{min}$;

According to [8], [9], $t$ is formally defined as in Eq. S9 for individuals with general cardinality [10], where $\alpha$ is the probability to visit all the points in the space, l is the number of genes, $\mu$ the mutation rate and *n* the population size [11].

$$t = \frac{\ln(1-\alpha)}{\ln(1-min\{(1-\mu)^{lr}, \mu^{lr}\})} \qquad (S9)$$

Since Eq. S9 is U shaped with the global minimum in $\mu = 0.5$, it is possible to use an approximation:

$$t = \frac{\ln(1-\alpha)}{\ln(1-0.5^{l})} \qquad (S10)$$

Thus, the values are set as in Eq. S11

$$\begin{cases} t_{min} = 400 & \alpha = 0.9 \\ t_{max} = 5500 & \alpha \to 1 \end{cases} \qquad (S11)$$

#### *SI 4: Formulation of the conditional probability for computing the Shannon entropy*

The derivation of the probability for evaluating the Shannon entropy (H) starts from the assumption that GA, modeled as a stochastic dynamical system, is Markovian: the actual state depends on the previous state and not others. This allows modelling the systems using a Markov Chain formalism and to simplify the derivation.

The probability to have the individual *i* at the generation j ($X_{ij}$) depends on the possible transformation ($T$) of its parent (s) *k*, belonging to the previous generation $X_{k(j-1)}$. Formally:

$$P(X_{ij}|X_{k(j-1)}, T) = \frac{P(X_{ij}, X_{k(j-1)}, T)}{P(X_{k(j-1)}, T)} \qquad (S12)$$

Note that since there are only crossover ($T_c$) and mutation ($T_m$) transformations, $T = T_c = T_m = 0.5$, and $k$ could correspond to a pair or a single parent(s), respectively. Moreover, in the crossover the event is modeled with a binomial distribution while in the mutation a uniform distribution is used: the events are equiprobable and the joint probability collapses to the marginal one.

Given the independency between the individual $X_{k(j-1)}$ and its transformation, the previous formula becomes:

$$P\left(X_{ij} \middle| X_{k(j-1)}, T\right) = \frac{P\left(X_{ij}, X_{k(j-1)}, T\right)}{P\left(X_{k(j-1)}, T\right)} = \frac{\left(P\left(X_{ij}\right) P\left(X_{k(j-1)}\right) P(T)\right)}{P\left(X_{k(j-1)}\right) P(T)} = P\left(X_{ij}\right) \quad \text{(S13)}$$

In this way, all the individuals are considered independent events of the dynamical system. Specifically, $X_{ij}$ is modeled as a $(1 \times 6)$ raw vector of the six morphological genes and his Shannon entropy $(H(X))$ is calculated using the Eq. S14.

$$\begin{cases} H(X) = -p(X) \log_2 p(X) \quad X \in \mathcal{R}^{+}, \mathbb{N} \quad (S14) \\ H(i) = \sum_{z=1}^{6} H(X_z) \qquad\qquad (S15) \\ H(g) = \sum_{i=1}^{50} H(i) \end{cases}$$

From this, the individual complexity $(H(i))$ and the total complexity of the generation $H(g)$ are determined (Eq. S15).

***SI 5 Hardware and Finite element computation***

All computations were performed on a PC/Intel core i9-10940X CPU with 3.30 GHz clock frequency and 64 GB RAM, running Windows 10 (64bit) operating system.

Determining the biophysical genes in the loop of evolution requires computation of the FF to score individuals across generations. Using the *Transport of Diluted Species* module in COMSOL, FE models are numerically solved in COMSOL at the steady state using the UMFPACK direct solver with a *finer* mesh size; a skewness factor between 0.5 and 0.8 is applied to maintain the mesh as regular as possible.

***Table SI 1:*** *Parameters used for FE simulations in Evolvoid.*

| Parameter | Value [unit] | Description |
|---|---|---|
| $D_{ox}$ | $3 * 10^{-9}$ $[m^2 * s^{-1}]$ | Diffusion coefficient of oxygen in water at 310.15 K [12] |
| $T_{cell}$ | $310.15$ $[K]$ | Environmental temperature |
| $ocr$ | $4.8 * 10^{-17}$ $[mol * s^{-1} cell^{-1}]$ | Average single-cell oxygen consumption rate in the construct[3] |
| $k_m$ | $7.39 * 10^{-3}$ $[mol * m^{-3}]$ | Michaelis-Menten constant of hepatic cells [3] |
| $\rho_{cell}$ | $5.14 * 10^{14}$ $[1 * m^{-3}]$ | Physiological cell density of the hepatic tissue [13] |
| $c_{0.2}$ | $0.2$ $[mol * m^{-3}]$ | Constant oxygen concentration boundary condition[14] |
| $c_{0.15}$ | $0.15$ $[mol * m^{-3}]$ | Constant oxygen concentration boundary condition for nutrient perturbation |

***SI 6: Fitness function (FF)***

Given the convexity of the FF function, a global maximum is ensured. Due to the normalization of the FF parameters (i.e., their adimensionality), the searching process is constrained in the working range [0,1] highlighted in Figure SI1.

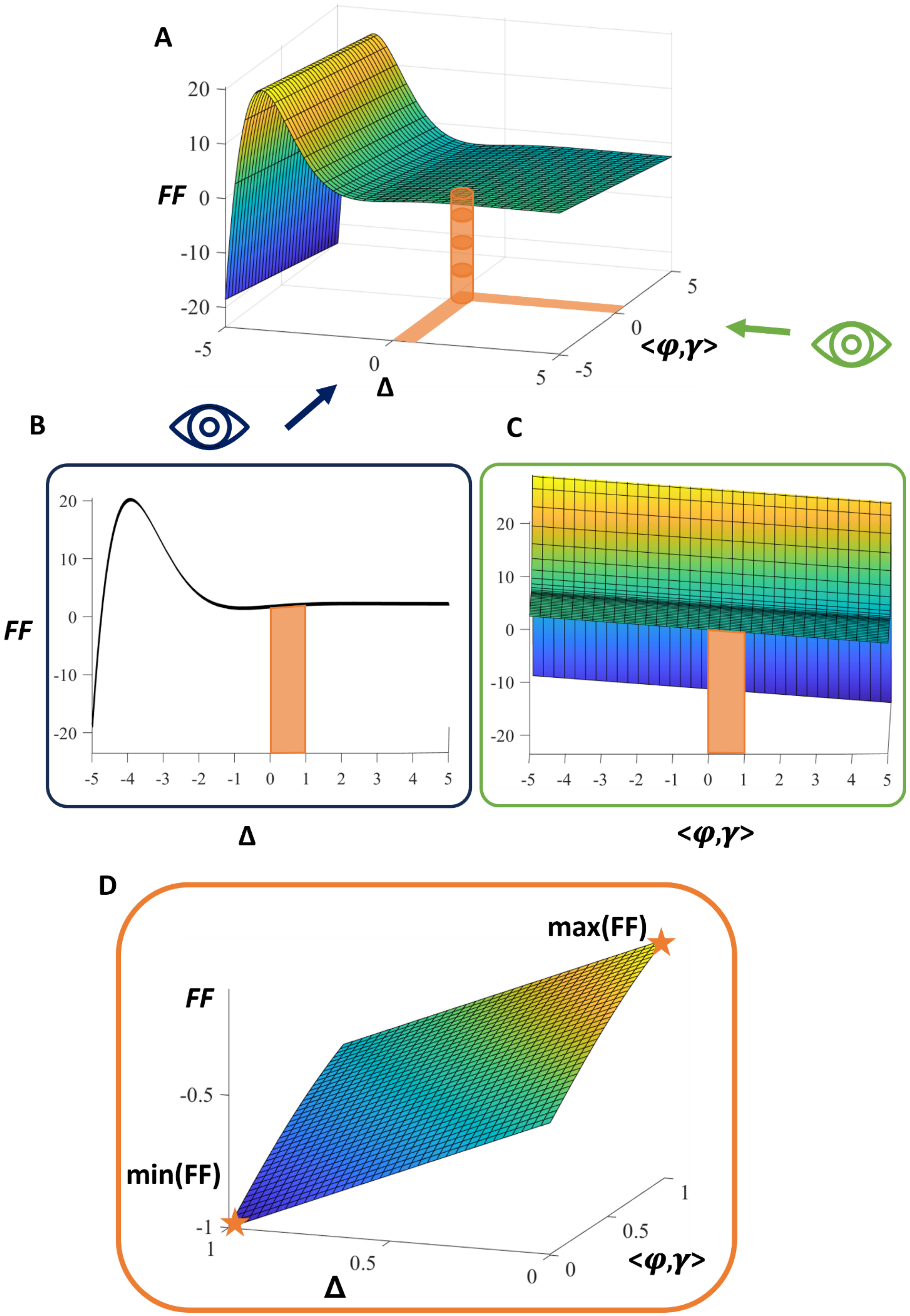
A
FF
20
10
0
-10
-20
-5
0
5
Δ
-5
0
5
<φ,γ>
B
FF
20
10
0
-10
-20
-5 -4 -3 -2 -1 0 1 2 3 4 5
Δ
C
20
10
0
-10
-20
-5 -4 -3 -2 -1 0 1 2 3 4 5
<φ,γ>
D
max(FF)
FF
-0.5
min(FF)
-1
1
0.5
0
Δ
1
0.5
0
<φ,γ>

***Fig. SI* 1.** *A visualization of the optimum search space driven by the FF from different points of view. A) plot of the FF as a function of* $\Delta$ *and* $< \varphi, \gamma >$*. B) and C) FF projection along* $\Delta$ *and* $< \varphi, \gamma >$*, respectively. D) The Evolvoid working space.*

*SI 6.a Analysis of the fitness function sensitivity to weights*

A sensitivity analysis was conducted in order to obtain the combination of $w_{\Delta}$ and $w_{<\varphi,\gamma>}$ complying with the following criteria: i) minimum number of GA cycle to the convergence according to assembly theory [15], ii) surface tension comparable to that measured in *ex vivo* embryonic tissues [16], [17], for obtaining physically and biologically relevant cellular models; iii) the last generation has to have a dispersion of the FF point as small as possible. The whole construct volume ($V$) is chosen as the morphological index to inspect for sensitivity, since it indirectly allows assessing for all the morphological parameters. The sensitivity of the FF to biophysical genes is obtained by analyzing both the sphericity ($\varphi$) and three-dimensionality ($\gamma$) with respect to the weights.

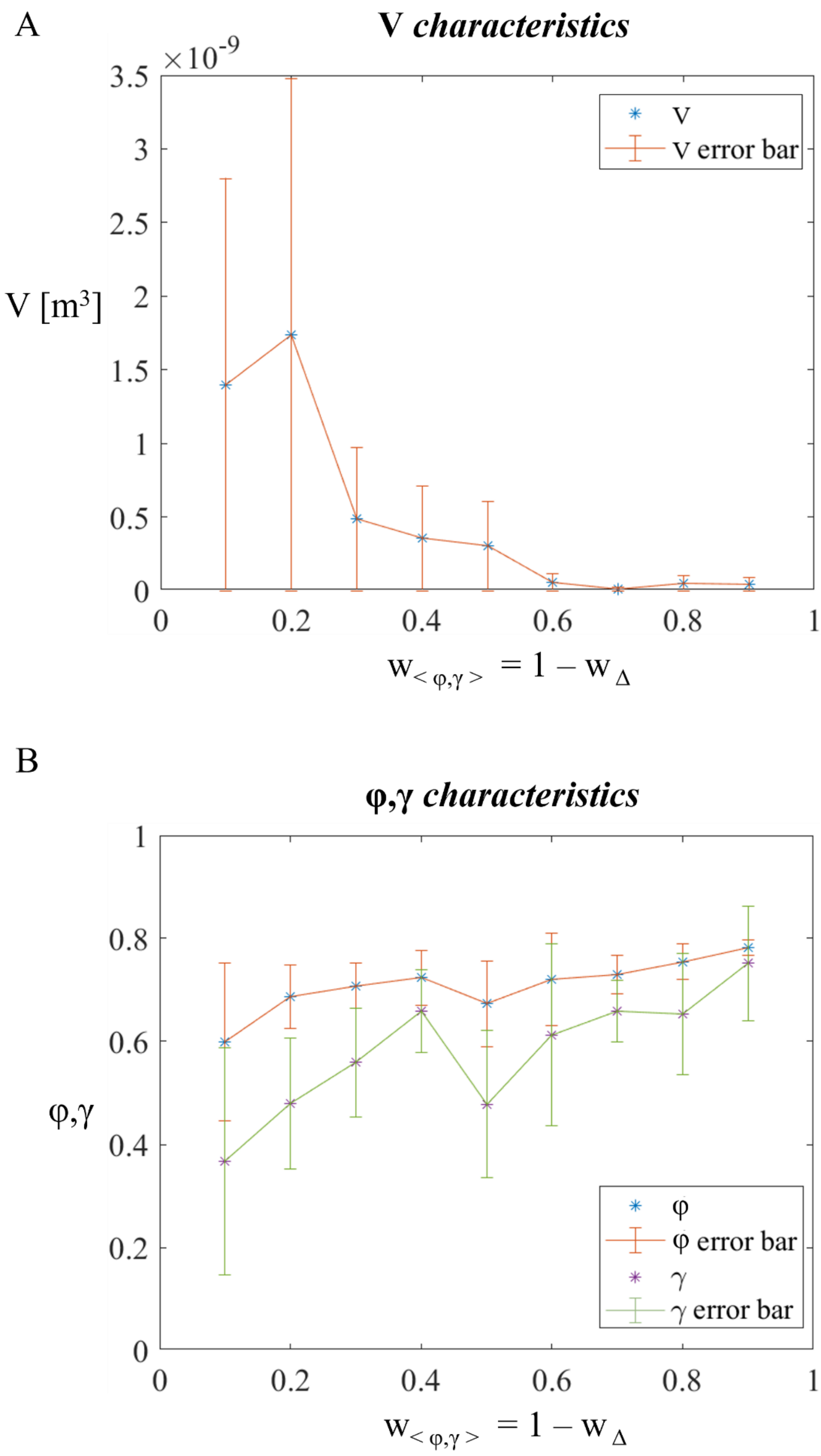


***Fig. SI2*** *A) Mean and standard deviation of constructs volume measured in ($m^3$) as a function of weights. B) Mean and standard deviation of constructs sphericity (φ) and three-dimensionality (γ) as a function of weights.*

Figure SI 2A show that the volume decreases with increasing $< \varphi, \gamma >$ associated weight: the algorithm constantly balances viability (Δ), shape $(\varphi, \gamma)$ and dimensions $(V)$ in order to maintain and $O_2$ diffusion characteristic time ($t_D$) less than the $O_2$ reaction characteristics time ($t_R$) (Eq. S16)

$$\begin{cases} t_D = \frac{L^2}{D} \\ t_R = \frac{1}{K} = \frac{k_m + c^*}{OCR * \rho} \end{cases} \tag{S16}$$

where $L$ is the characteristic length ($r$), $c^*$ the maximum oxygen concentration value for considering the worst case scenario and all the others values as in Table SI 1. Since all the parameters are constrained, the only way to assure the condition $t_D < \tau_R$ is to decrease the dimension of the construct.

On the other hand, the other characteristics (Fig. SI 2B) demonstrate the tendency of $\varphi$ and $\gamma$ to increase with increase of the $< \varphi, \gamma >$ associated weight. In conclusion, the sensitivity analysis shows that the combination of weights $w_\Delta = 0.5$ and $w_{<\varphi,\gamma>} = 0.5$ represent the optimal balance for the three characteristics.

***SI 7: Successive generation***

Crossover allows densifying the search space, since it generates individuals owning differently combined but already existing morphological features; on the other hand, random mutation increases the diversity of the population, enabling the exploration of a wider spectrum of possible solutions.

Properly balancing the densification and expansion of the search space across generations is crucial to avoid suboptimal solutions (*i.e.,* local rather than global maxima of the *FF*). In this light, the fraction of non-elite parents undergoing crossover is randomly set according to a gamma probability density [18], and the fraction of mutated progeny is consequently derived as specified in Table SI2. The population to be evaluated at each iteration of the loop is thus constituted by the offspring, in turn composed as follows:

$$offspring = elite + crossover + mutation \tag{SI17}$$

Once the population is completed, the loop of evolution is iteratively run until a stopping criterion is verified (*SI 3).*

***Table SI2****. Fractions of the population of the i-th generation determining the offspring (i.e., the population of the (i+1)-th generation) genome through different mechanisms.*

| ***Parameter*** | ***Value*** | ***Description*** |
|---|---|---|
| *Elite fraction (E)* | 0.05 | Elitist individuals correspond to the best-scored 5% of the population. |
| *Gene recombination fraction* | $1 - E$ | Fraction of non-elitist individuals undergoing gene recombination. |
| *Crossover fraction (C)* | $\Gamma(\gamma = 4.94)$ | The fraction of non-elitist individuals undergoing crossover is picked from a gamma probability distribution [18]. |
| *Mutation fraction (M)* | $1 - E - C$ | The fraction of individuals undergoing mutation is set as the remaining non-elitist individuals |

### *References*


[1] O. Roeva, S. Fidanova, and M. Paprzycki, ‘Influence of the Population Size on the Genetic Algorithm Performance in Case of Cultivation Process Modelling’.

[2] MathWorks, ‘Global Optimization Toolbox User’s Guide R2023b’, 2004. [Online]. Available: www.mathworks.com

[3] A. Ahluwalia, ‘Allometric scaling in-vitro OPEN’, 2017, doi: 10.1038/srep42113.

[4] E. Botte, F. Biagini, C. Magliaro, A. Rinaldo, A. Maritan, and A. Ahluwalia, ‘Scaling of joint mass and metabolism fluctuations in in silico cell-laden spheroids’, *BIOPHYSICS AND COMPUTATIONAL BIOLOGY*, 2021, doi: 10.1073/pnas.2025211118/-/DCSupplemental.

[5] D. Xu *et al.*, ‘The ellipsoidal area ratio: an alternative anisotropy index for diffusion tensor imaging’, *Magn Reson Imaging*, vol. 27, no. 3, pp. 311–323, Apr. 2009, doi: 10.1016/J.MRI.2008.07.018.

[6] F. J. Rohlf, ‘Morphometrics’, *Annu Rev Ecol Syst*, vol. 21, no. 1, pp. 299–316, 1990, doi: 10.1146/ANNUREV.ES.21.110190.001503.

[7] M. S. Steinberg, ‘Reconstruction of Tissues by Dissociated Cells’, *Science (1979)*, vol. 141, no. 3579, pp. 401–408, Aug. 1963, doi: 10.1126/science.141.3579.401.

[8] H. Aytug and G. J. Koehler, ‘Stopping Criteria for Finite Length Genetic Algorithms’, *https://doi.org/10.1287/ijoc.8.2.183*, vol. 8, no. 2, pp. 183–191, May 1996, doi: 10.1287/IJOC.8.2.183.

[9] H. Aytug and G. J. Koehler, ‘New stopping criterion for genetic algorithms’, *Eur J Oper Res*, vol. 126, no. 3, pp. 662–674, Nov. 2000, doi: 10.1016/S0377-2217(99)00321-5.

[10] G. J. Koehler, S. Bhattacharyya, and M. D. Vose, ‘General Cardinality Genetic Algorithms’.

[11] A. E. Nix and M. D. Vose, ‘Modeling genetic algorithms with Markov chains’, *Ann Math Artif Intell*, vol. 5, no. 1, pp. 79–88, Mar. 1992, doi: 10.1007/BF01530781/METRICS.

[12] C. Magliaro, A. Rinaldo, and A. Ahluwalia, ‘Allometric Scaling of physiologically-relevant organoids’, *Scientific Reports 2019 9:1*, vol. 9, no. 1, pp. 1–8, Aug. 2019, doi: 10.1038/s41598-019-48347-2.

[13] E. Bianconi *et al.*, ‘An estimation of the number of cells in the human body’, *Ann Hum Biol*, vol. 40, no. 6, pp. 463–471, Nov. 2013, doi: 10.3109/03014460.2013.807878.

[14] A. Carreau, B. El Hafny-Rahbi, A. Matejuk, C. Grillon, and C. Kieda, ‘Why is the partial oxygen pressure of human tissues a crucial parameter? Small molecules and hypoxia’, *J Cell Mol Med*, vol. 15, no. 6, pp. 1239–1253, Jun. 2011, doi: 10.1111/J.1582-4934.2011.01258.X.

[15] A. Sharma, D. Czégel, M. Lachmann, C. P. Kempes, S. I. Walker, and L. Cronin, ‘Assembly theory explains and quantifies selection and evolution’, *Nature* |, vol. 622, 2023, doi: 10.1038/s41586-023-06600-9.

[16] G. Forgacs, R. A. Foty, Y. Shafrir, and M. S. Steinberg, ‘Viscoelastic Properties of Living Embryonic Tissues: a Quantitative Study’, *Biophys J*, vol. 74, pp. 2227–2234, 1998, doi: 10.1016/S0006-3495(98)77932-9.

[17] R. A. Foty, ’, G. Forgacs, C. M. Pfleger, and M. S. Steinberg’, ‘Liquid Properties of Embryonic Tissues: Measurement of Interfacial Tensions’, vol. 72.

[18] M. S. Mcpeek and T. P. Speed, ‘Modeling Interference in Genetic Recombination’, 1995, Accessed: Dec. 01, 2023. [Online]. Available: https://academic.oup.com/genetics/article/139/2/1031/6013227
